# Direct visualization of native infectious SARS-CoV-2 and its inactivation forms using high resolution Atomic Force Microscopy

**DOI:** 10.1101/2020.10.23.351916

**Authors:** Sébastien Lyonnais, Mathilde Hénaut, Aymeric Neyret, Peggy Merida, Chantal Cazevieille, Nathalie Gros, Christine Chable-Bessia, Delphine Muriaux

**Affiliations:** CEMIPAI, University of Montpellier, UMS3725 CNRS, Montpellier, France; Institute of Research in Infectiology of Montpellier (IRIM), University of Montpellier, UMR9004 CNRS, Montpellier, France; Université de Montpellier, Institut des Neurosciences de Montpellier (INM), Montpellier, France

## Abstract

The severe acute respiratory syndrome coronavirus 2 (SARS-CoV-2) is responsible for COVID19, a new emerging pandemic affecting humans. Here, single viruses were analyze by atomic force microscopy (AFM) operating directly in a level 3 biosafety (BSL3) facility, which appeared as a fast and powerful method to assess infectious virus morphology in its native conformation, or upon inactivation treatments, at the nanoscale level and in 3D. High resolution AFM reveals structurally intact infectious and inactivated SARS-CoV-2 upon low concentration of formaldehyde treatment. This protocol allows the preparation of intact inactivated SARS-CoV-2 particles for safe use of samples out of level 3 laboratory, as revealed by combining AFM and plaque assays, to accelerate researches against the COVID-19 pandemic. Overall, we illustrate how adapted BSL3-atomic force microscopy is a remarkable toolbox for rapid and direct virus identification and characterization.

## Main

SARS-CoV-2 has emerged as a new zoonotic coronavirus in late 2019 in Wuhan, China, and rapidly spread worldwide, causing a global pandemic ^1,2^. This novel virus is classified as a hazard group 3 pathogen in most countries, which thus requires researches in level 3 biosafety facilities. Proper inactivation methods are therefore crucial to transfer the material from high confinement space towards a standard laboratory, facilitating researches under safe working conditions. Virus inactivation can be achieved in many ways, like the application of heat, alcohol, peroxide, radiation, fixatives or detergents, and have been described for coronaviruses^3–6^. However, these procedures may impair the structural features of the viruses, which can be necessary for many applications and diagnosis based on morphology. Published experimental data that demonstrate SARS-CoV-2 inactivation keeping intact particles are lacking. We therefore proposed here a simple method allowing the inactivation of SARS-CoV2 virions while maintaining their native morphology.

We first controlled that the virus isolate was SARS-CoV-2 by means of revealing the presence of the M, N and S viral proteins in infected cell lysates using immunoblots, by quantitative RT-PCR, targeting the E gene, overtime on the cell culture supernatant and by imaging fixed infected cells by Transmission Electron Microscopy (TEM) (supplementary Figure 1). One can see that the infected cells are producing SARS-CoV-2. We then directly visualized native infectious SARS-CoV-2 from the clarified infected cell supernatant using a BSL3 customized AFM, which is a non-destructive and non-intrusive imaging technique particularly powerful to directly visualize biological samples in physiological conditions, such as viruses^7–9^. This AFM major advantage is its simple physical operating principle^10^, which allows a fast 3D analysis of hydrated, intact particles and virion size distribution in buffer, without any treatment or staining, after smooth adsorption of the sample on a poly-L-lysine coated mica surface. We implemented this instrument in BSL-3, which allowed us to image native infectious SARS-CoV-2 particles at high resolution in quantitative imaging AFM mode within minutes from cell supernatant (Figure 1a) or after a simple washing procedure (Figure 2). Native infectious SARS-CoV-2 virions appeared on the AFM images as roughly spherical or ellipsoidal (Figure 1), consistent with electron micrographs^11,12^. Surprisingly, most of the viral particles appeared embedded in a network of thin filamentous structures adhering to the surface (Figure 1a and supplementary Figure 2), in addition to free particles, suggesting that SARS-CoV-2 particles are released form the infected culture cells in packages. As noted elsewhere^12^, spikes of the S protein appear highly fragile and the majority of AFM images showed “bald” viral particles without or very low S trimers protruding from the viral surface, even from viruses directly imaged from cell supernatant. This is probably due to virion movements on the mica surface during adsorption, causing them to lose and/or break the long spike proteins. We found however some viruses studded with thin, nail-like structures ranging from 14 to 22 nm, attached to the virion membrane and flattened on the mica surface, suggesting Spike trimers, as in ^11^ (Figure 1b). Using height profiles in AFM, particle diameters ranged from 45 to 140 nm, with mean diameter of 91 nm, consistent with diameters measured by cryo-electron tomography for SARS-CoV-2 ^11,12^ and for other coronaviruses^13^. TEM observation of the virus producing cells confirmed unambiguously the typical structure of coronavirus particles, containing granular densities corresponding to the viral nucleoprotein and showing protruding spikes S proteins incorporated into the viral lipid envelope^14^ (Figure 1D). By TEM, dehydrated particles show diameters distribution in the range from 40 to 95 nm, in good accordance with the known ≈25% shrinkage of samples with the dehydration procedure used here for sample preparation^15^.

**Figure 1.**
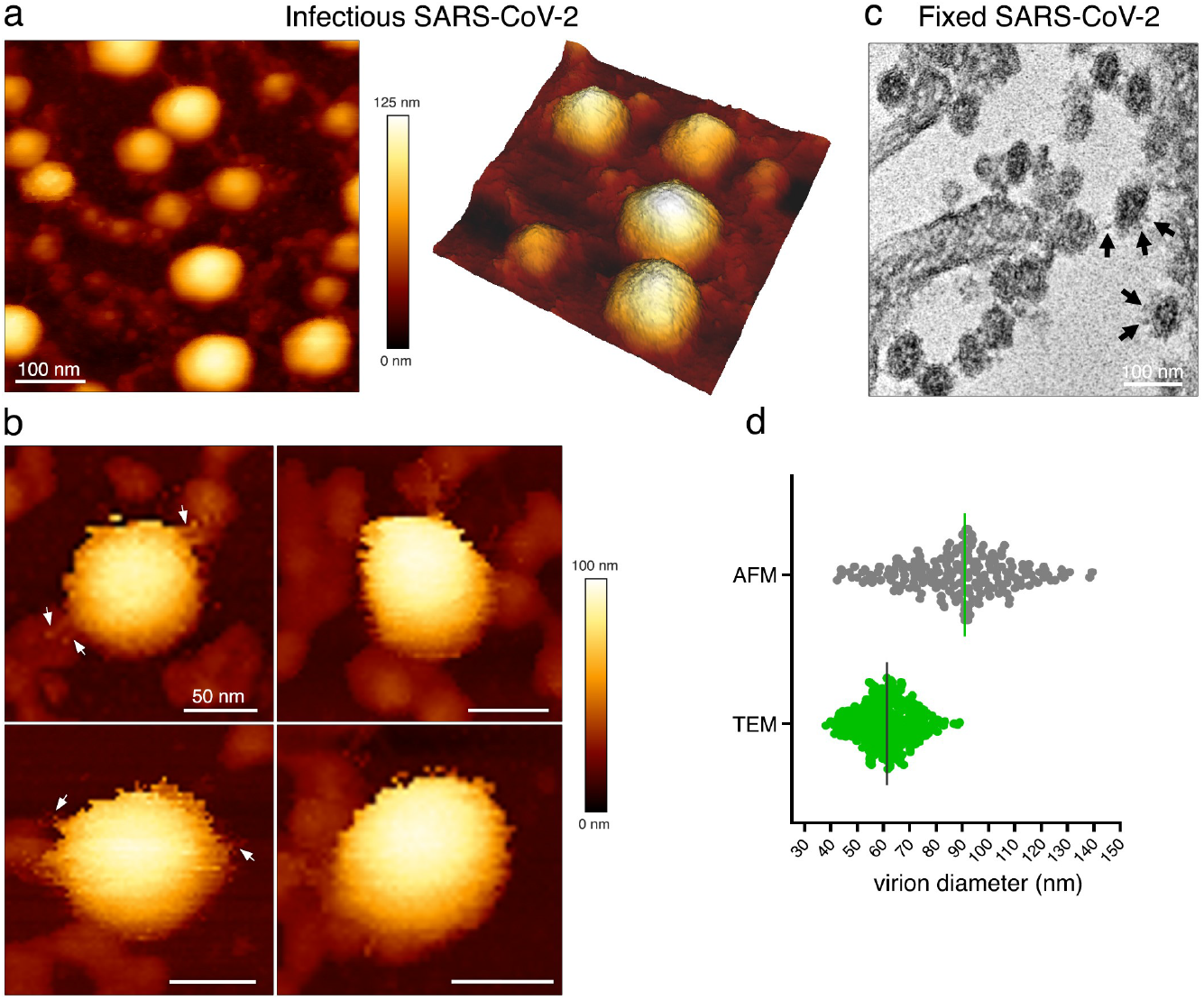
Native infectious SARS-CoV-2 virions imaged by high resolution BSL3-AFM. **(a)** Topographic image and 3D projection of native infectious SARS-CoV-2 virions adsorbed on a mica surface using quantitative imaging (QI) mode AFM in buffer. (**b**) Zoom-in view of SARS-CoV-2 virions studded with nail-like structures (arrows) ranging from 14 to 22 nm suggesting S-proteins (scale bars: 50 nm). (**c**) Example of a TEM image of fixed infected cells producing SARS-CoV-2 that can be seen as spherical particles studded with S trimers (see arrows). (**d**) Virion diameter distribution of infectious SARS-CoV2 samples imaged by AFM in liquid and compared to fixed samples imaged by TEM.

**Figure 2.**
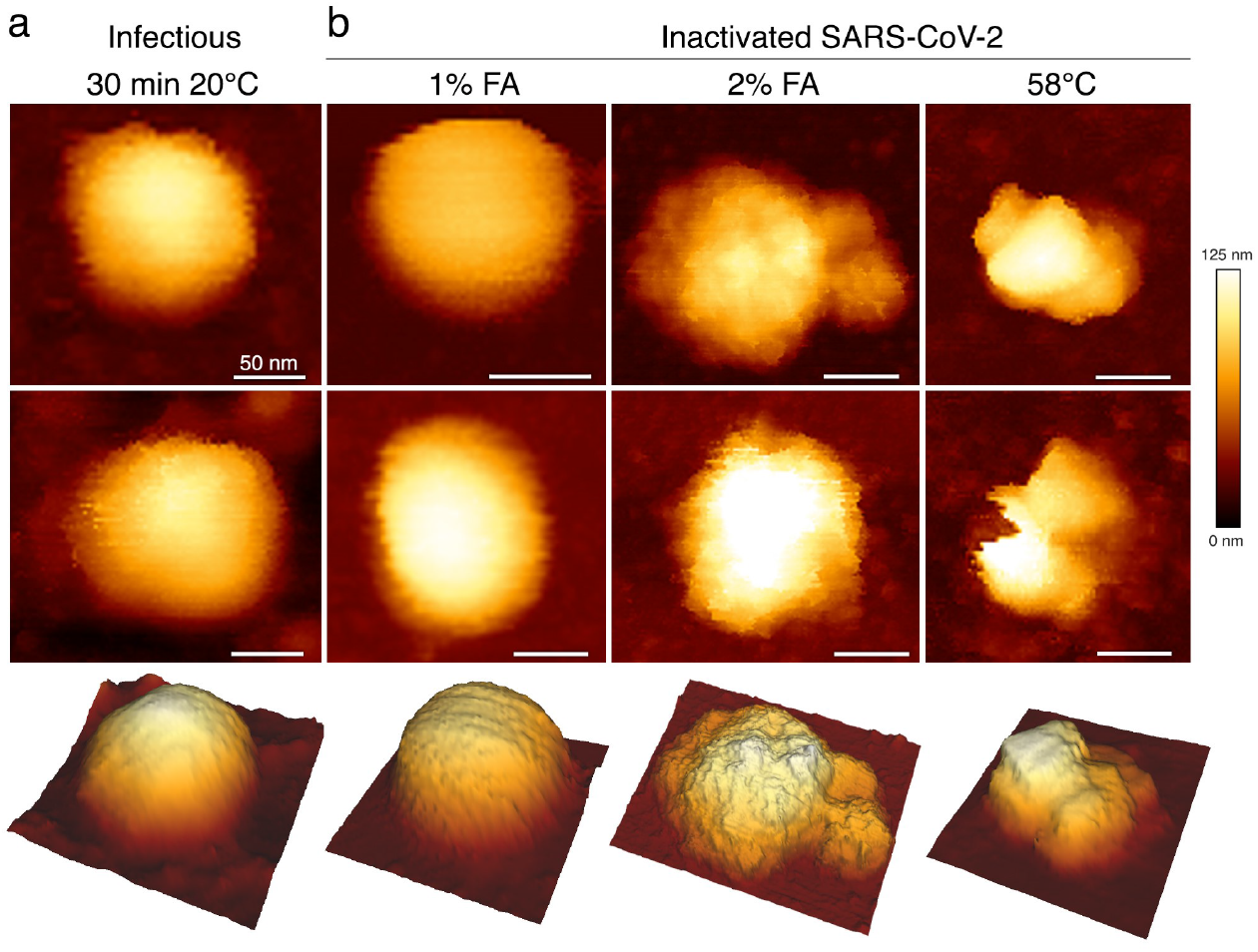
SARS-CoV-2 inactivation monitored by topographic imaging using QI mode AFM in buffer. **a**, Native infectious, control sample kept for 30min at room temperature. **b**, Virus particles inactivated with 1% and 2% FA for 30min at 20°C, or incubated at 58°C for 30min. Scale bars are 50 nm. The color gradient for Z scale is the same for all AFM panels. Two representative images and a 3D projection are shown in column for each virion.

We then examined and compared virus inactivation using formaldehyde (FA), and by heat, to identify a condition keeping structurally intact SARS-CoV-2 particles. Upon heat or FA treatments, followed by ultrafiltration, viral particles were analyzed directly by AFM for morphology (compare Figure 2a for WT with no treatment with Figure 2b with treatments) and plaque assays for infectivity (Table 1, supplementary Figure 3). SARS-CoV-2 incubated at 58°C for 30 minutes was successfully inactivated (Table 1), as described for other coronavirus^4,5^. AFM analysis showed severely damaged particles that have lost their spherical shapes (Figure 2b). Aldehyde fixation is well established in histology and EM^16^. Unlike glutaraldehyde, which can cause aggregates, buffered FA has been shown to preserve virus morphology^17^, and its virucidal efficiency is very well documented, e.g., in vaccination programs^18,19^. We observed complete inactivation of SARS-CoV-2 after incubation at 20°C for 30 min with 0.5%, 1%, 2% or 3.6% FA (see Table 1) evaluating infectivity by plaque assays (supplementary Figure 3). AFM analysis showed unaltered particles with a shape similar to the untreated control upon 0.5% or 1% FA treatment (Figure 2b), while the higher percentage dramatically damaged the viruses, whose morphology is then similar to the samples heated to 58 °C (Figure 2b). As shown previously^4,17^, FA inactivation effect was temperature dependent and was strongly reduced at 4°C. Indeed, SARS-CoV-2 was still infectious for a lower concentration of 0.1% FA with an incubation at 4°C (Table 1, supplementary Figure 3). Altogether, these results show that SARS-CoV-2 at a concentration of 1-2.10^6^ PFU/ml are inactivated at 20°C using 0.5% or 1% FA for 30min at RT, without loss of virus physical integrity. In contrario, keeping the virus at 4°C in buffer, or up to 48h at 20°C, has no incidence on its titer (Table 1) indicating a strong stability of this virus in buffer, probably due to its embedded filamentous form (Figure 1a and supplementary Figure 2).

**Table 1:**
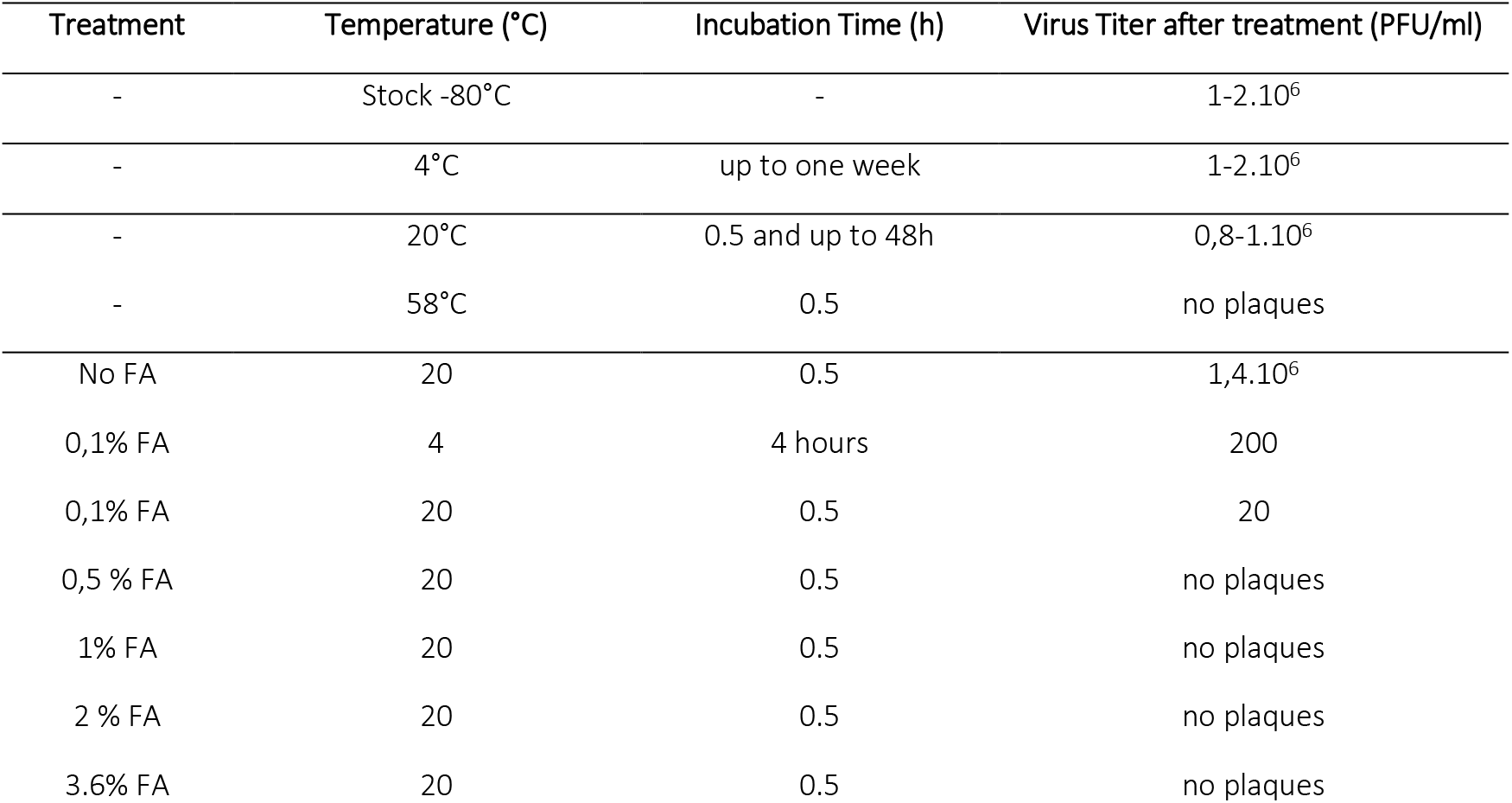
Efficiency of SARS-CoV-2 inactivation by FA in relation to temperature and incubation time. Virus titer was determined by plaque assays (see Supplementary Figure 2).

| Treatment | Temperature (°C) | Incubation Time (h) | Virus Titer after treatment (PFU/ml) |
| --- | --- | --- | --- |
| - | Stock -80°C | - | 1-2.10<sup>6</sup> |
| - | 4°C | up to one week | 1-2.10<sup>6</sup> |
| - | 20°C | 0.5 and up to 48h | 0,8-1.10<sup>6</sup> |
| - | 58°C | 0.5 | no plaques |
| No FA | 20 | 0.5 | 1,4.10<sup>6</sup> |
| 0,1% FA | 4 | 4 hours | 200 |
| 0,1% FA | 20 | 0.5 | 20 |
| 0,5 % FA | 20 | 0.5 | no plaques |
| 1% FA | 20 | 0.5 | no plaques |
| 2 % FA | 20 | 0.5 | no plaques |
| 3.6% FA | 20 | 0.5 | no plaques |

In summary, in front of the urgent need to perform researches on SARS-CoV-2 to fight COVID-19, the inactivation methods described here will contribute to transfer the virus from the confined laboratory to lower biosafety class laboratory in its inactivated form preserving its particle round shape. We show that low percentage of FA treatment allow to retain the physical integrity of the particles albeit non infectious. High resolution AFM analysis of SARS-CoV-2 in buffer also demonstrates its ability to provide direct and fast qualitative information on infectious virus morphology and proves to be a method of choice for analyzing viral preparations with exceptional precision and rapidity.

## Materials and Methods

### Virus and cell culture

VeroE6 cells (African green monkey kidney cells) were obtained from ECACC and maintained in Dulbecco’s minimal essential medium (DMEM) supplemented with 10% heat inactivated fetal bovine serum (FBS) at 37°C with 5% CO_2_. The strain BetaCoV/France/IDF0372/2020, was supplied by the National Reference Center for Respiratory Viruses hosted by Institut Pasteur (Paris, France) and headed by Pr. Sylvie van der Werf. The human sample from which strain BetaCoV/France/IDF0372/2020 was isolated has been provided by Dr. X. Lescure and Pr. Y. Yazdanpanah from the Bichat Hospital, Paris, France. Moreover, the BetaCoV/France/IDF0372/2020 strain was supplied through the European Virus Archive goes Global (EVAg) platform, a project that has received funding from the European Union’s Horizon 2020 research and innovation program under the grant agreement No 653316. For supplementary Figure 3 (panel d) we used a SARS-CoV-2 strain isolated from CPP Ile de France III, n°2020-A00935-34 and CRB Collection of the University Hospital of Montpellier, France (www.chu-montpellier.fr). Both virus strains were propagated in VeroE6 cells with DMEM containing 2.5% FBS at 37°C with 5% CO_2_ and were harvested 72 hours post inoculation. Virus stocks were stored at −80°C.

### Quantitative reverse transcription polymerase chain reaction (qRT-PCR)

RNAs from mock infected or infected (MOI=0.001) cell culture supernatant were extracted using the Nucleospin Dx Virus RNA purification kit (Macherey-Nagel). Then qRT-PCR was performed in triplicate as described^20^, using primers targeting the E gene of SARS-CoV-2 (E_Sarbeco-F ACAGGTACGTTAATAGTTAATAGCGT; E_Sarbeco-R ATATTGCAGCAGTACGCACACA) and Luna Universal One-Step qRT-PCR Kit (New England Biolabs) on a Roche Light Cycler 480. The calibration of the assay was performed with a nCoV-E-Sarbeco-Control Plasmid (Eurofins Genomics).

### Western Blot

VeroE6 cells were infected for 2h with SARS-CoV-2 (MOI = 0.001). 96h post infection, cells were washed twice in PBS, detached with versen, pelleted at 250 x g for 6min and lysed in RIPA buffer. Total protein concentration was calculated using a Bradford protein assay kit. 20μg of total cell lysates were diluted in Laemmli buffer and proteins were separated by SDS-PAGE on 8% (for S-protein) and 10% (for M- and N-protein) acrylamide gels. Gels were transferred to PVDF membrane using wet transfer with Tris-glycine-methanol buffer. Membranes were washed in TBS, blocked with 5% milk in TBS-T for 30min and incubated overnight at 4°C with primary antibody specific for spike (Gentex, cat# GTX632604), N-protein (Gentex, cat# GTX632269) or M-protein (Tebu, cat# 039100-401-A55), all three diluted at 1:1,000 in TBS-T. After washing with 5% milk in TBS-T, the membranes were incubated with HRP conjugated anti-mouse antibodies for N and S protein, and with HRP conjugated anti-rabbit antibody for M protein for 2h at room temperature, washed again in TBS-T, incubated with ECL reagent (Amersham cat#RPN2236) and imaged using a Chemidoc Imager (Biorad).

### Virus inactivation

All inactivation conditions were performed with a starting viral stock of 1-2.10^6^ plaque forming units/ml (PFU/ml). Heat inactivation was performed by incubating 100 μl of SARS-CoV-2 stock at 58°C for 30 min. Formaldehyde inactivation were performed by incubating 90μl of virus supplied by 10x FA-Hepes (0.5M) at 4°C or RT (20°C) for 30min, 2h or up to overnight at 4°C depending on the conditions described in Table 1. Samples of Figure 2 and supplementary Figure 3 (panels a,b,c “column washed”) were complemented with PBS up to 5 ml and washed using centrifugal concentrator (Microsep advance 100KDa MWCO, Pall corporation) at 1000xg at 4°C for 5 minutes. Samples were washed 4 times by filling again the concentrator with 5ml of PBS and repeating the centrifugation. The last centrifugation was processed until 200 μl remained, the virus sample was collected and stored at 4°C for one night before plaque assays. Samples of supplementary Figure 3 (panel d “ultracentrifuged”) were diluted into 4ml PBS after FA inactivation, loaded over a 20% sucrose cushion-TNE and ultracentrifuged at 100.000xg for 1h on a SW55ti Beckman rotor. Viral samples were resuspended in 20μl DMEM and stored at −80°C. Virus inactivation were monitored in plaque assay on a monolayer of VeroE6 cells (3,5.10^5^ cells/well), using 200 μl of virus solution. Samples were serially diluted and the PFU/mL values were determined using crystal violet coloration and subsequent scoring the amounts of wells displaying cytopathic effects.

### Atomic Force Microcopy

Freshly cleaved muscovite mica sheets (V1 grade, Ted Pella, Inc.) were glued on glass slide, coated for 10 minutes at 20°C with 0.1% poly-L-lysine (Sigma), rinsed with 3 ml of buffer A (10 mM Tris–HCl pH 7.5 and NaCl 100 mM) and dried with a N2 flux. A plastic O-ring (JPK instruments) was then glued on the glass slide to assemble a small liquid cell. Virus samples were diluted four-fold in buffer A and 200 μL were deposited on the mica for 15 minutes to allow passive virus adsorption. The liquid cell was next completed with 200 μl of buffer A before imaging. AFM imaging was performed at room temperature on a NanoWizard IV atomic force microscope (JPK Instruments-Bruker, Berlin, Germany) mounted on an inverted optical microscope (Nikon Instruments, Japan) and operating in a BSL3 laboratory. Topographic imaging was performed in quantitative imaging (QI) mode, which is a force-curve based imaging mode, using BL-AC40TS-C2 cantilevers (mean cantilever spring constant k_cant_ = 0.09 N/m, Olympus). Before each experiment, sensitivity and spring constant (thermal noise method) of cantilever were calibrated. The applied force was kept at 0.15 nN and a constant approach/retract speed of 40 μm.s-1 (z range of 100 nm). Images were flattened with a polynomial/histogram line-fit with the JPK data processing software. Low-pass Gaussian and/or median filtering was applied to remove minor noise from the images. The Z-color scale in all images is given as relative after processing, with the offset being kept the same within each of the figures to emphasize the structural features.

### Electron Microcopy

VeroE6 cells were infected with 1.10^6^ PFU of SARS-CoV-2 for 24h. Cells were fixed with 2,5% (v/v) glutaraldehyde in PHEM buffer and post fixed in osmium tetroxide 1% / K_4_Fe(CN)_6_ 0,8%, at room temperature for 1h for each treatment. The samples were then dehydrated in successive ethanol bathes (50/70/90/100%) and infiltrated with propylene oxide/ EMbed812 mixes before embedding. 70 nm ultrathin cuts were made on a PTXL ultramicrotome (RMC, France), stained with OTE/lead citrate and observed on a Tecnai G2 F20 (200kV, FEG) TEM at the Electron Microscopy Facility MRI-COMET, INM, Plate-Forme Montpellier RIO Imaging, Biocampus, Montpellier.

## Acknowledgements

We thank Dr Olivier Moncorgé, Dr Caroline Goujon and Dr Raphaël Gaudin from The Institut de Recherche en Infectiologie de Montpellier (IRIM) for initial advices in virus production and plaque assay setup. We are grateful to Dr Edouard Tuaillon and Dr Vincent Foulongne for provision of the SARS-CoV-2 strain from the Centre de Ressources Biologiques collection of the University Hospital of Montpellier, France, and to Dr Monsef Benkirane (IGF, Montpellier, France) for providing the first amplification of this virus on VeroE6 cells. This study was supported by the CNRS and Montpellier University through a Montpellier Université d’Excellence (MUSE) support. The BSL3-AFM was funded by the REDSAIM project.

## Author contributions

NG, MH and CCB, performed viral stock amplification, purification and titer; MH did viral infection, plaque assays and calculate titer; DM and SL set up and performed virus inactivation protocols, SL performed AFM imaging and analysis; AN, TEM sample preparation; AN and CC performed TEM imaging; PM performed infectious cell lysates and immunoblots; PM, AN, and SL setup and performed qRT-PCR; SL and DM edit the figures and wrote the manuscript, DM raise funding and direct the study.

## Supporting Information

**Supplementary Figure 1.**
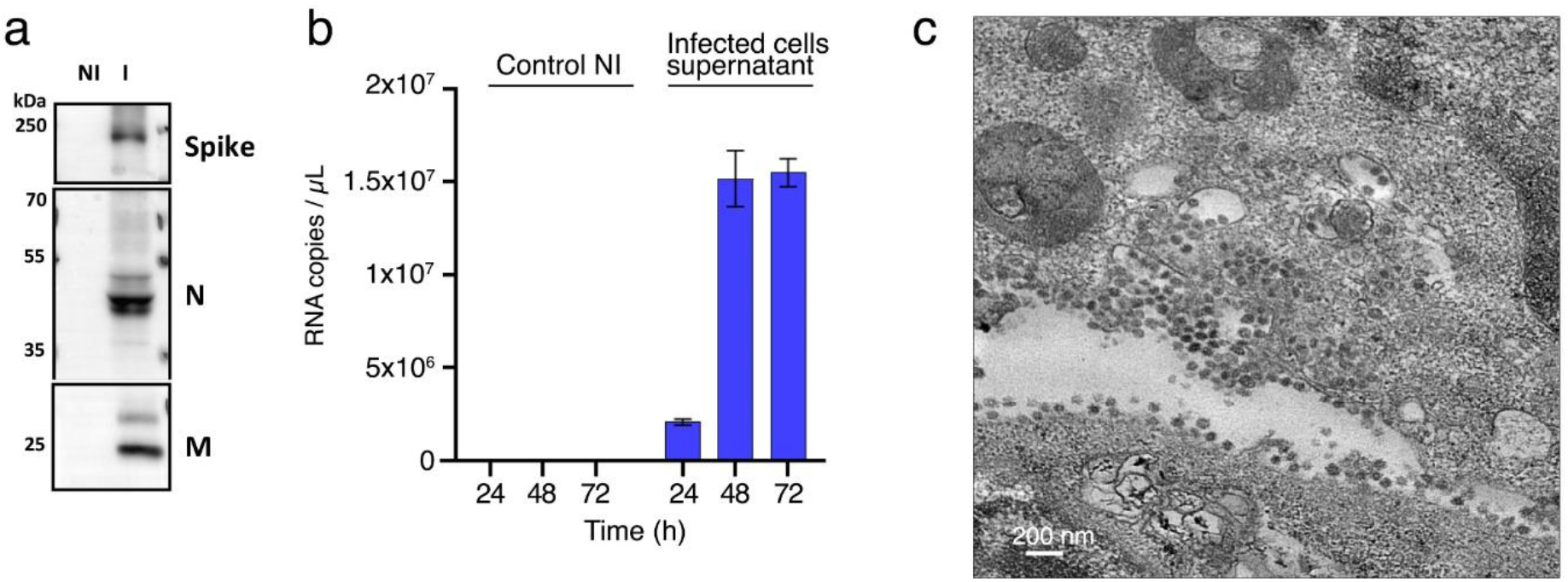
SARS-CoV-2 identification. (a) Western Blot analysis of SARS-CoV-2 S, M and N proteins in lysates of infected (I) and non infected (NI) VeroE6 cells. (b) SARS-CoV-2 production in the cell supernatant followed by qRT-PCR targeting the E gene. (c) An example of TEM image of SARS-CoV-2 infected VeroE6 cells for 24h (MOI=0.1).

**Supplementary Figure 2.**
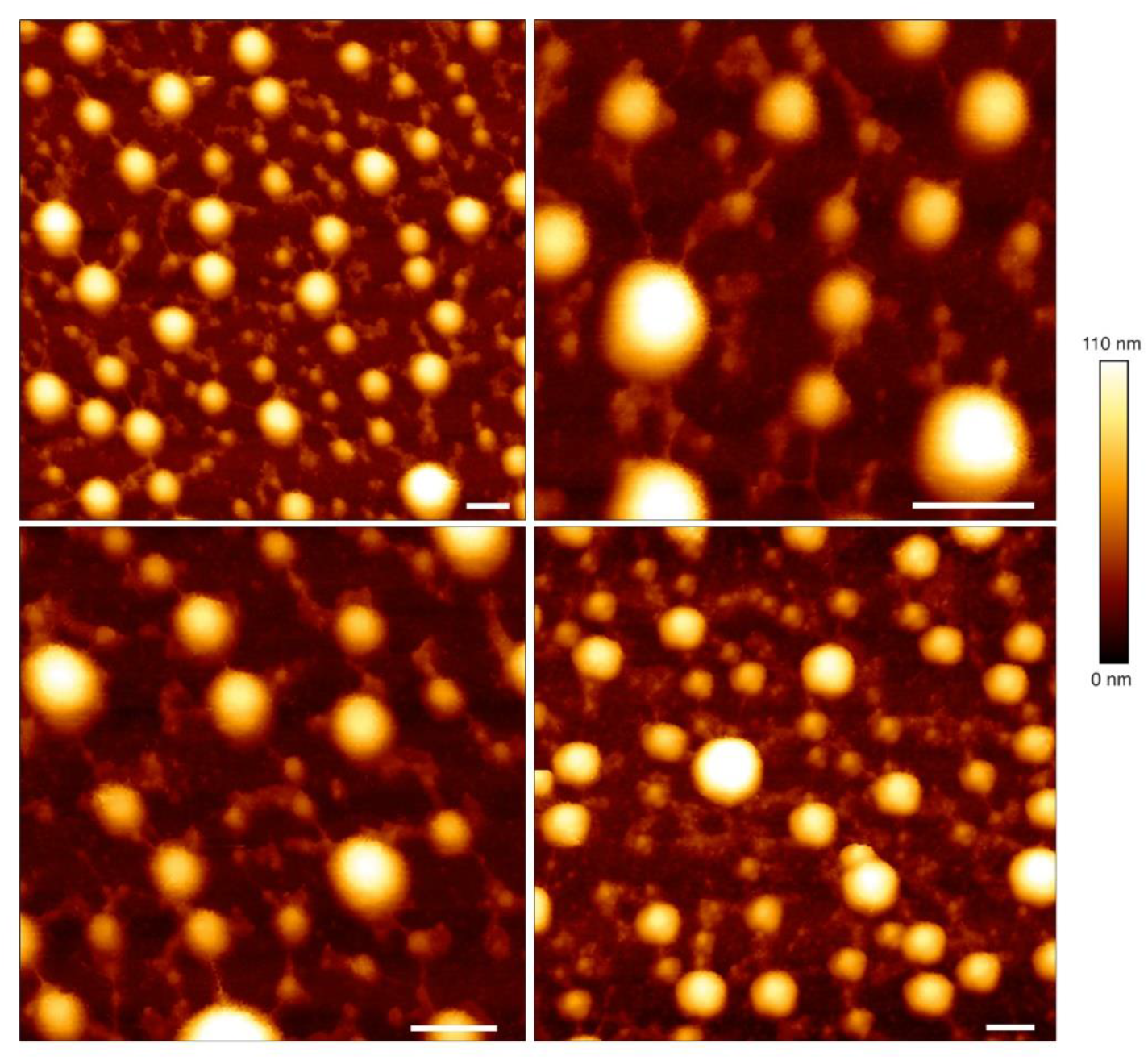
AFM images of native infectious SARS-CoV-2 found embedded in a network of filamentous structures. AFM topographic images in QI mode of infectious native SARS-CoV-2 from cell supernatant directly adsorbed on a mica surface using QI mode AFM in buffer. Scale bars: 200 nm.

**Supplementary Figure 3:**
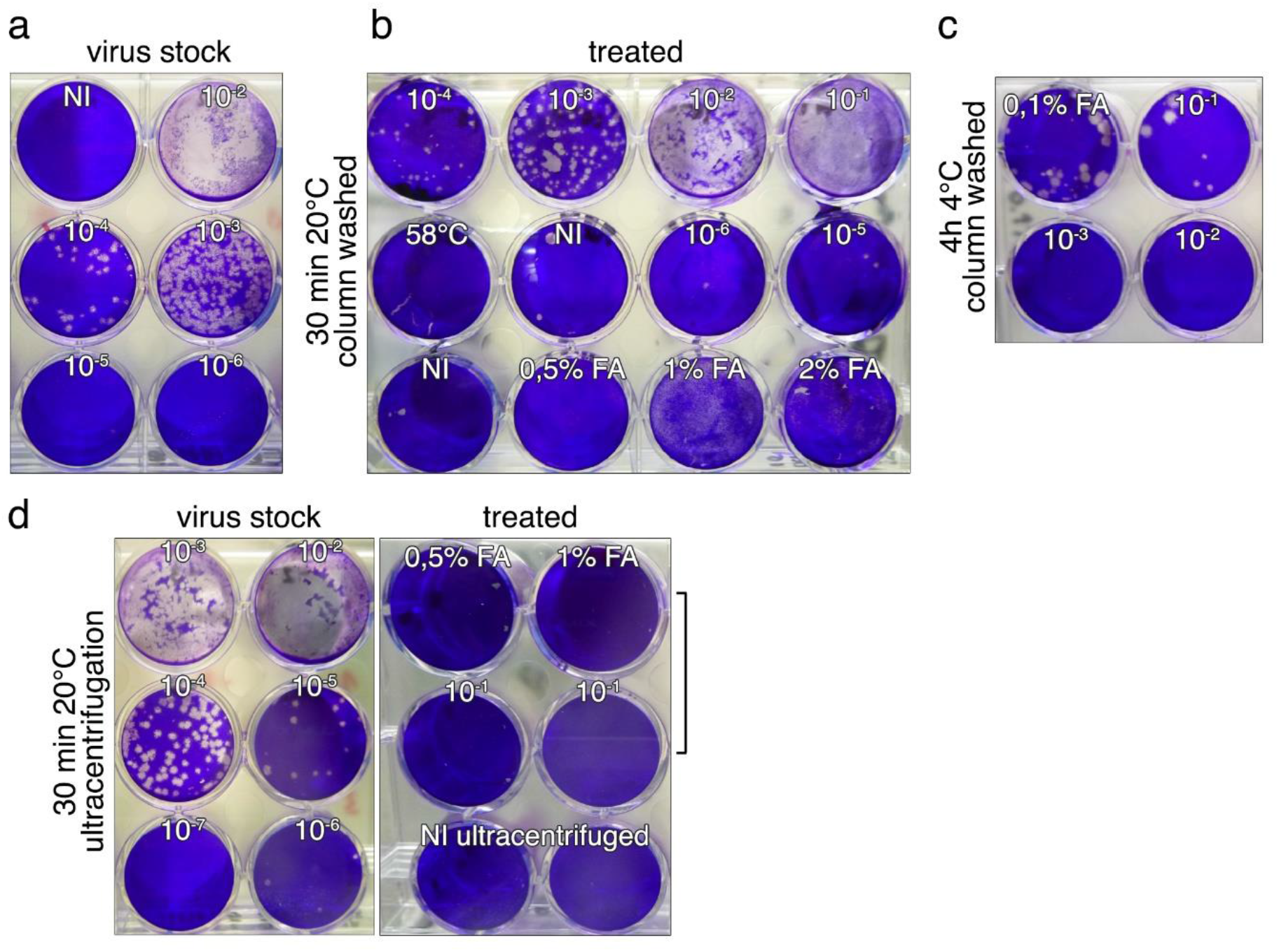
Native and inactivated SARS-CoV-2 titration. (a) SARS-CoV-2 titration using plaque assays (NI: non infected cells). (b) 0.1% to 2% FA and temperature inactivation of the virus analyzed by plaque assays.

